# Sleep/wake synchronization across latitude

**DOI:** 10.1101/378737

**Authors:** José María Martín-Olalla

**Affiliations:** Departamento de Física de la Materia Condensada. Universidad de Sevilla. Ap Correos 1065, ES41080. Sevilla

**Keywords:** sleep/wake cycle, sleep onset, time use survey, circadian rhythm, homeostatic sleep pressure, dst, daylight saving time, summer time

## Abstract

Analysis of time use surveys in seventeen European countries and two American countries suggest that the winter sunrise —the latest sunrise year round— is a synchronizer for the sleep/wake cycle in standard population below 54° latitude, in competition with the noon synchronizer.

When comparing industrialized data to data from hunter/gatherer, pre-industrial, Tropical societies only the late event survives as a synchronizer below 54° latitude. People rise immediately before sunrise —winter sunrise in industrialized mid-latitude societies— and abhor morning darkness. Synchronization propagates through the sleep/wake cycle so that people go to bed with increasing distance to sunset in winter as latitude increases in a scenario dominated by artificial light. This suggests a leading role of the homeostatic sleep pressure in understanding sleep/wake cycle at social level.

**WARNING:** *This pre-print has been largely upgraded and restyled in* “Sleep timing in industrial and pre-industrial societies sync to the light/dark cycle” *(https://doi.org/10.1101/392035). In this new pre-print data coming from the Harmonsized European Time Use Surveys (HETUS) and referred to the “sleep/wake and other personal care” cycle were not analyzed. Instead two new pre-industrial data are included.*

*Therefore this old pre-print you are about to read remains as a source for these data (see Figure 3 and Table III, Table VII to IX) and a source of information in the range of latitude above 54°.*

## I INTRODUCTION

In the last three decades sleep research has addressed basic questions related to sleep/wake activity: what triggers sleep, what triggers awakening[1; 2] or why do we sleep late and struggle to wake up on time[3]. The two process model of sleep regulation[4] defines a framework in which a homeostatic process (Process S), which depends on previous wake times, and a process controlled by the circadian pacemaker (Process C), which do not depend on previous wake times, regulate sleep onset.

It is also a day-to-day evidence that sleep/wake cycle is synchronized to Earth’s rotation period T = 24 h —the definition of hour— and entrained to the light/dark cycle.[5] This cycle is also shaped by Earth’s obliquity *ε* = 23.5°—the angle between Earth’s rotation axis and Earth’s orbital axis— which gives rise to seasons and seasonality —the adjustment of human activity through seasons. Understanding the role of latitude[6; 7; 8; 9; 10; 11] in human cycles is an open issue.

Recently Yetish *et al.*[12] reported natural sleep seasonal variations in three pre-industrial, tropical societies. They noted pre-industrial societies typically awaken before sunrise and go to sleep several hours after sunset. Also recently Martin-Olalla[13] reported evidences of the role of the winter day as a synchronizer of human basic activities — sleeping, working, eating, among others— in industrialized, modern, Western societies, including sleep-wake cycle in employees.

This work extends this analysis to standard population (twenty to seventy years old, employees and non-employees, weekday and weekend) and compares industrialized societies with pre-industrial data from Yetish *et al.*[12]. Winter sunrise, the latest sunrise time year round, j will be confirmed as a synchronizer for human sleep/wake cycle below 54° latitude. That applies to both risetimes and bedtimes which suggests a predominant role of homeostatic sleep pressure (Process S) in sleep regulation at a social level.

## II METHODS

National time use surveys (NTUS, hereafter) from six European countries and two American countries will be analyzed to characterize the sleep-wake cycle in standard population (twenty to seventy four years old). Survey microdata were either publicly available[14; 15; 16] or could be obtained through petitions[17; 18; 19; 20; 21] and are those of Martin-Olalla[13], which analyzed sleep/wake cycle in employees.

Sleep-wake cycle data from hunter/gatherer-horticulturalist pre-industrial societies were retrieved from Yetish *et al.*[12] (Table S2). They include the San people, in Kalahari, the Hadza people in the most equatorial area of Tanzania, and the Tsimane people in Bolivia, all of them in the Antarctic hemisphere and within the tropical range. While NTUS data accounts for some 40 % of the population living in their latitudinal range, data of pre-industrial societies are much less relevant in this sense.

On the second hand the “sleep and other personal care” cycle in standard population obtained from NTUS and from the pre-prepared tables available at the Harmonized European Time Use Survey[22] (hereafter, HETUS) website will be analyzed. HETUS is a merger of fifteen European time use surveys prior to 2005. Sleep-wake cycle and “sleep and other personal care” cycle slight differ in a small lag in the morning: people usually wake up —changing status in the sleep-wake cycle— and do some personal care like grooming or dressing. Only when the other personal care ends, the status is changed in the “sleep and other personal care” cycle. At bedtime, a second, smaller lag is also observed.

The analysis will contrast sleep/wake human activity against the seasonal light/dark cycle, which depends on latitude. Table I lists geographical data of countries to be analyzed in this work. Data represent population weighted median values retrieved from the database of cities with a population larger than 1000 inhabitants at http://www.geonames.org. Ephemerides were built up from them. Table I also lists values corresponding to the pre-industrial societies.

Sleep/wake and “sleep and other personal care” cycles can be statistically analyzed by their daily rhythms: the shares of population which are doing the activity as a function of time within one cycle (day). Except for the lag both rhythms are similar: they roughly look like a smoothed rectangular function (see Figure 1 in Martin-Olalla[13]) with eventually some distortions due to naps. Rectangular shaped rhythm is a primary evidence of human collective behavior: if people were randomly sleeping or awaken during one cycle the daily rhythm should roughly yield a constant value. To some extend this behavior may be driven socially but their bulk properties show up the circadian rhythm and its entrainment to the light/dark cycle.

**Figure 1.**
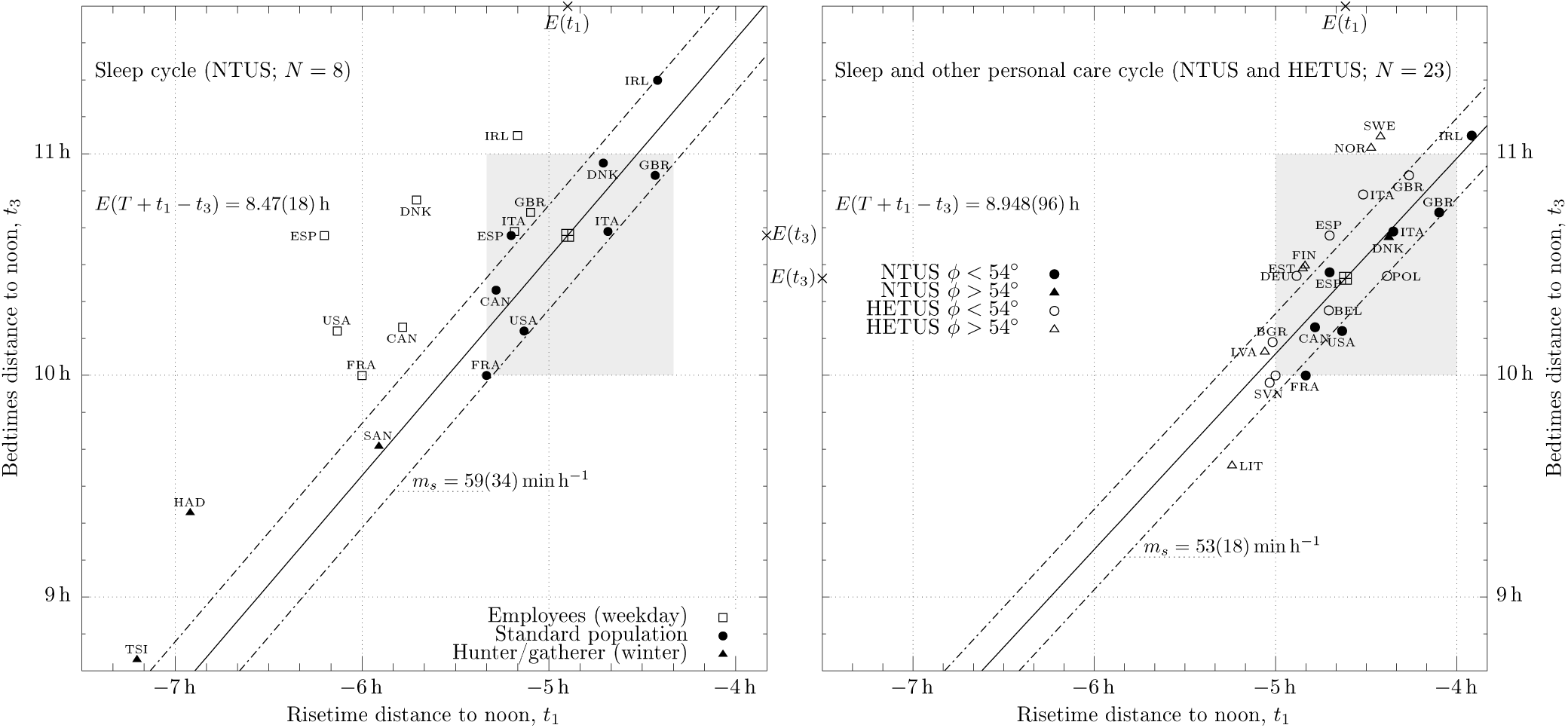
Bedtimes distance to noon *t*_3_ vs risetimes distance to noon *t*_1_ during winter time. Left panel show data from the sleep/wake cycle for standard population (solid squares), employees in a weekday (open squares) and hunter/gatherer pre-industrial societies from Yetish *et al.*[12]. Right panel shows data from the “sleep and other personal care cycle” referred to standard population. Straight line fits standard population data; on right panel it only accounts for countries below *φ* = 54*°*. The crossed box displays distribution average values, also marked on complementary axes. Dash-dotted lines show the norm of the residuals squared below and above the fit result. Panels show slope (confidence interval at the standard level) and average value of *t*_3_ *- t*_1_. Country labels display iso-3166-1 alpha-3 codes. Data are listed in Table II and Table III.

A threshold located at half the distance from maximum to minimum daily rhythm helps defining two relevant timemarks in the daily rhythm assessing when the daily rhythm overshoots (risetime, *t*_1_) and undershoots (bedtime, *t*_3_) the threshold. A third time mark can be measured by assessing when half of the daily activity has been consumed and half remains. This timemark is slightly different from the mid point of risetime and bedtime. It will be termed hereafter *wakeful noon time, t*_2_.

Every of these timemarks can be expressed in different ways. Raw microdata provide local time values but they are unuseful for comparing to the light/dark cycle. Instead natural gauges like time distance to solar noon, time distance to sunrise/sunset, or, eventually, solar elevation angle at timemarks are the appropriate measures for testing against the light/dark cycle.

The winter photoperiod *D*_*w*_ —the shortest photoperiod year round— will be the predictor in this analysis. It is a function of latitude *φ*:

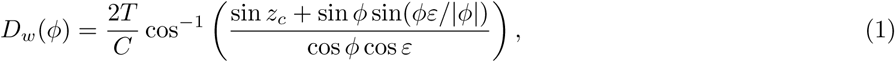

where *C* = 360° is one cycle and *z*_*c*_ is a critical solar altitude relative to horizon which defines sunrise and sunset. Equation (1) gently simplifies and some symmetries appear for *z*_*c*_ = 0°; however a more accurate value, which takes into account solar angular size and atmospheric refraction, is *z*_*c*_ = −0.83°. Notice that *D*_*w*_(*φ*) is only defined and non-zero below polar circles.

Winter photoperiod is a convenient proxy for latitude because it draws linear relations for the most significant events of the light/dark cycle. Winter sunrise and sunset can simply be cast as:

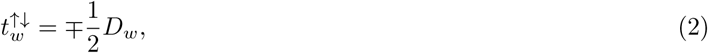

where time is measured relative to solar noon. If *z*_*c*_ is set to 0° the relation exactly holds in summer when signs are reversed and time is measured relative to midnight. Therefore *β* = 1*/*2 = 30 min h^-1^ characterizes the gradient of the boundaries of the light/dark cycle: it reads thirty minute in advance/delay of winter/summer sunrise/sunset per hour change in winter photoperiod. Gradient is negative (advance) for the “late events” —winter sunrise or summer sunset— and positive (delay) for the “early events” —winter sunset or summer sunrise. Gradient *β* must be contrasted to the gradient of “mean events” —solar noon, mean sunrise and mean sunset times— which is exactly zero.

Summarizing, three human-related timemarks —risetimes, bedtimes, wakeful noon times— for standard population will be contrasted against *D*_*w*_ to test which light and dark event —late (*m* = *-β*), noon/mean (*m* = 0) and early (*m* = *β*)— best explains synchronization.

Notice a competition between modern mechanical clocks, which are synced to noon, and ancient, pre-industrial clocks, which are related to sunrise and sunset, is implicit in the analysis.

### III RESULTS

Table II lists sleep/wake cycle mean annual timemarks from NTUS data from hunter/gatherer, pre-industrial societies[12]. Table III lists “sleep and other personal care” timemarks obtained from NTUS and HETUS.

Figure 1 shows on the left panel data from the sleep/wake cycle. For comparison NTUS data from employees sleep-wake cycle in a weekday[13] are also shown. For the latest set risetimes are located one hour earlier than standard population while bedtimes mostly coincide. Average difference *E*(*T* +*t*_1_ *-t*_3_) approximately resembles daily sleep time and amounts to 8.47 h for standard population. It is some forty minutes smaller for employees (*E*_emp_(*T* + *t*_1_ *- t*_3_) = 7.8 h) and some twenty-five minute smaller for pre-industrial societies in winter (*E*_hg_(*T* + *t*_1_ *- t*_3_) = 8.1 h).

Figure 1 shows in right panel data from the “sleep and other personal care” cycle. Average difference is slightly larger, which is understandable since it casts two activities in one rhythm.

Both panels fit standard population data into a one hour width box located ten to eleven hours after noon (10pm to 11pm solar time) and five to four hours before noon (7am to 8am solar time) on the right panel; risetimes are twenty minutes earlier on the left panel. One hour is a standard for human time variability, linked to one-hour width geographical time zones and to our preference for whole clock hours.

Data exhibit a positive linear relationship: earlier risetimes display earlier bedtimes. They are close to a one-to-one relation: slopes are *m* = 59(34) min h^-1^ (standard population, left panel) and *m* = 53(18) min h^-1^ (countries below 54°, right panel), with values in parenthesis showing confidence interval at the standard level *α* = 5 %.

Figure 2 displays mean annual risetimes, wakeful noon times and bedtimes from standard population sleep-wake cycle against shortest photoperiod and winter data from pre-industrial societies. Background colors highlight the photoperiod (lightest background) and night (darkest background) in the winter day. Dotted lines shows winter solar elevation angles starting at *z* = −12° (outermost lines, which defines nautical twilight) in steps of 6°.

**Figure 2.**
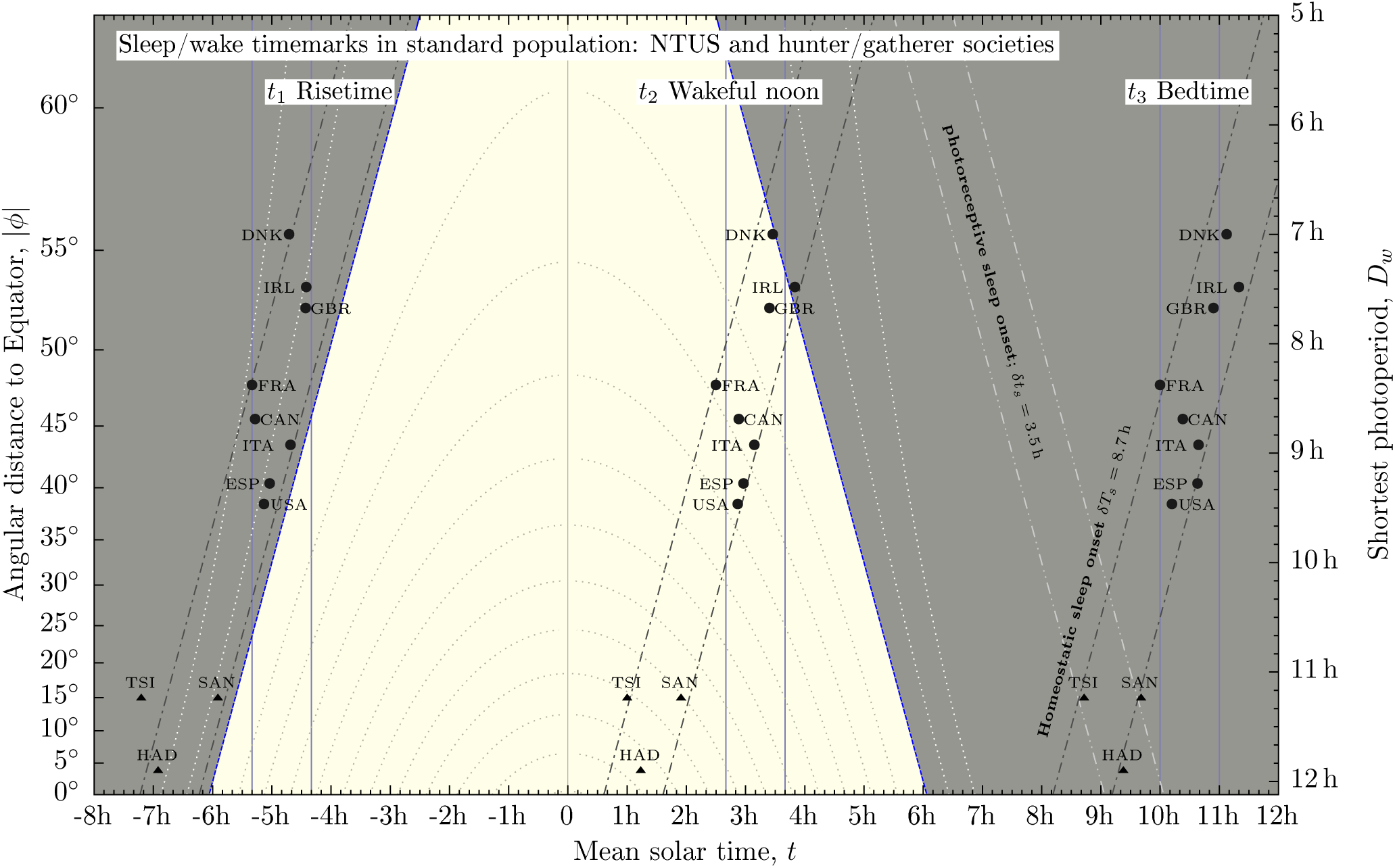
Timemarks extracted from standard population mean annual sleep-wake cycle in NTUS (circles) and hunter/gatherer pre-industrial societies from Table S2 in Yetish *et al.*[12] (triangles) against shortest photoperiod (right axis) or latitude (left axis). Blue solid lines show winter and summer terminator (*z* = −0.83°). Dotted lines show solar elevation angle starting at *z* = −12° (outermost lines) in steps of 6°. Lightest background shows winter photoperiod; darkest background is winter night. Vertical solid lines highlight distance to noon. Slanted dash-dotted lines highlight distance to late winter sunrise. Slanted dash-dot-dot lines highlight distance to winter sunset. Geographical data are listed in Table I; timemarks appear in Table II.

NTUS data can be placed into a vertical, one-hour width box highlighting distance to noon synchronization. Notwithstanding this they can also be placed into a one-hour width, slanted strip with slope *- β* (dash-dotted lines in the figure), which highlight distance to late event (winter sunrise or summer sunset) synchronization. Pre-industrial low-latitude hunter/gatherer data find a remarkable coincidence in this box.

To elucidate which event best explains statistical data variance multiple linear regression analyses were performed in GNU-octave —a free software mostly compatible with MatLab— by the function regress. Every regression analysis takes *D*_*w*_ as the predictor *x* —notice, however, it is displayed on the vertical axis in Figure 2. The response *y* is given by timemarks distance to noon —horizontal axis in the figure. Roughly speaking the idea is to ascertain whether points are randomly placed within vertical boxes or they do so within slanted strips.

This analysis will be summarized in three univariate descriptive properties of the response: range, sample variability and sample mean; and four bivariate descriptive properties: slope, confidence intervals at the standard level for slope and mean value, Pearsons’s coefficient *R*^2^ and *p*-value. The null hypothesis *H*_0_ in the analysis is “the response {*y*_*i*_} does not depend on the predictor {*x*_*i*_}”. Therefore a *p*-value smaller than the standard level *α* = 5 % suggests *H*_0_ can be statistically rejected at this level. The hard conclusion is the event unlikely synchronizes the timemark by alone. On the contrary, if *p > α*, the null hypothesis sustains at this level and the soft conclusion is the event may likely synchronize the timemark.

Results are listed in Table IV for distance to solar noon timemarks. Table V analyzes distance to winter sunrise and solar elevation angle. Finally Table VI inspects the scenario in summer.

A similar analysis can be done for the “sleep and other personal care” cycle. In this case NTUS data are accompanied HETUS data, which helps extending the latitudinal range poleward. Pre-industrial values are not available now. Figure 3 shows experimental data within the seasonal light/dark cycle: lightest background shows the region of permanent photoperiod, bounded by the winter terminator; darkest background shows the region of permanent dark, bounded by the summer terminator; intermediate background highlights the region where light and dark swings as seasons go by.

**Figure 3.**
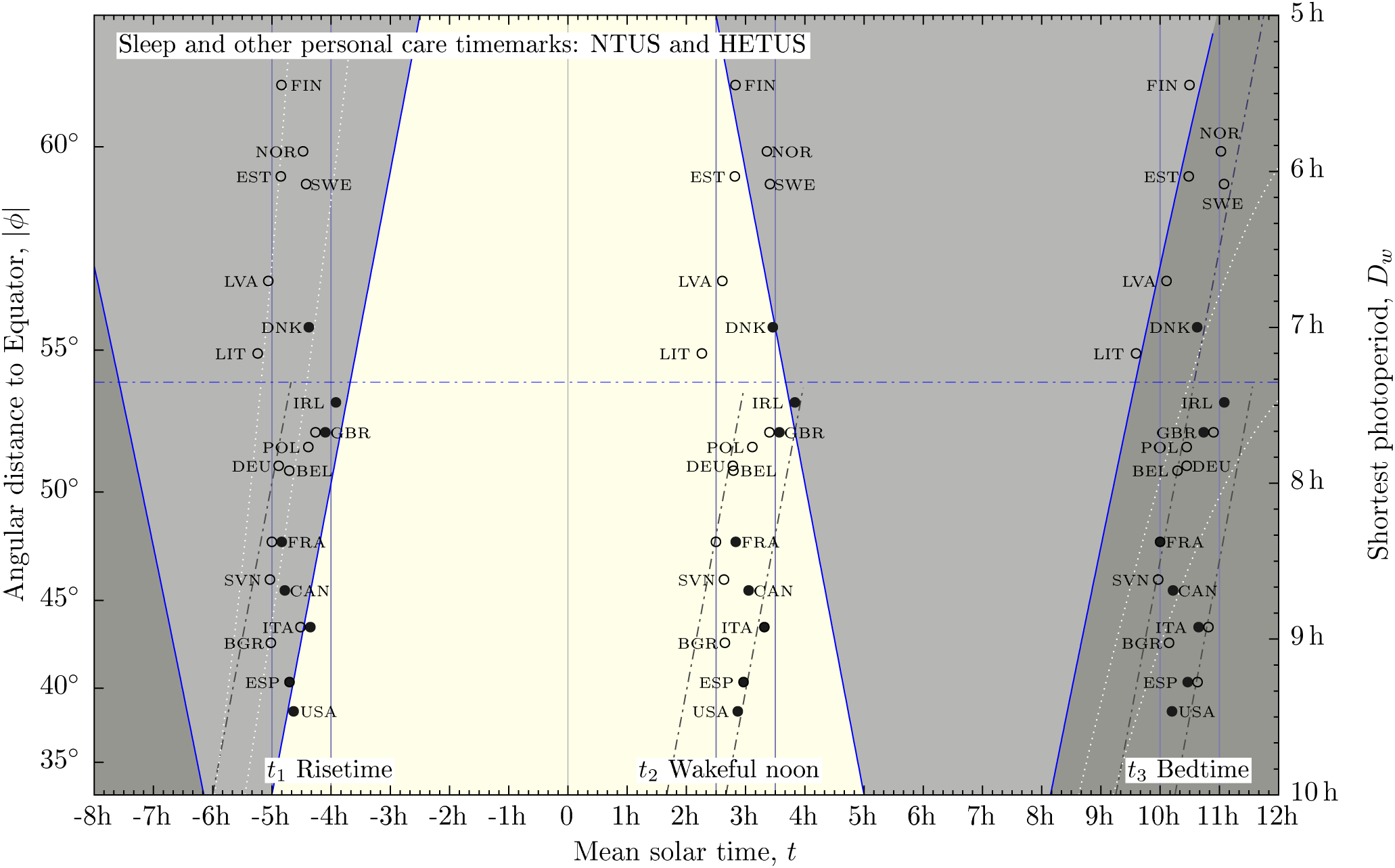
Timemarks extracted “sleep and other personal care” cycle against shortest photoperiod (right axis) or latitude (left axis). Bold circles show data from NTUS; open circles, data from HETUS. Lines and background colors highlight the cycle of light and dark. Lightest background is the region of permanent photoperiod; darkest background is the region of permanent dark; intermediate background shows the region that swings from light to dark as seasons go by. Blue solid lines show winter and summer terminator (*z* = −0.83°). Dotted lines the civil (*z* = −6°) and nautical (*z* = −12°) twilight lines in winter (dawn) and summer (dusk). Vertical solid lines highlight distance to noon. Slanted dash-dotted lines highlight distance to late event (winter sunrise and summer sunset). Summer lines have been delayed for one hour to accommodate daylight saving time. Geographical data are listed in Table I; timemarks appear in Table III.

Risetimes lie in the neighborhood of the winter sunrise. Data points can be placed in vertical, one-hour width strip for the full range of latitude. However data below 54° can be placed in slanted, one-hour width strips. That separates Scandinavian and Baltic data points from the rest of evidences.

Multiple linear regression analysis results are listed in Table VII (all data); Table VIII (latitudinal range 36*° < φ <* 54°) and Table IX (*φ >* 54°).

## IV DISCUSSION

### IV.I Sleep/wake cycle: Equator to mid-latitudes

The burden of this work is to ascertain the statistical relevance of correlations between sleep related times and shortest photoperiod (latitude). Figure 2 provides basic visual information which is always a key to start such analysis. Two major restrictions jeopardizes the analysis: sample size (*N* = 8) and the range of the predictor, which is slightly greater than two hours (see Table IV). That makes the range of winter sunrise time roughly one hour; the standard measure of human time variability. Therefore placing NTUS risetimes in vertical one-hour width boxes or in slanted one-hour width boxes requires further analysis.

Table IV show regression results for *N* = 8 timemarks related to the sleep-wake cycle. Risetimes hit a slope equal to *m*_*s*_ = −17(19) min h^-1^, close to *-β/*2. The parenthesis displays the confidence interval at the standard level *α* = 5 % which, in units of *β*, extends from −1.20 to 0.09. The confidence interval includes both late (*m* = *-β*) and noon (*m* = 0) events; on the contrary it excludes the early event *m* = +*β*. Therefore, (1) the null hypothesis “risetimes distance to noon do not depend on *D*_*w*_” sustains at this level: noon is a likely synchronizer for risetimes; (2) the null hypothesis “risetimes distance to winter sunrise do not depend on *D*_*w*_” also sustains at this level and winter sunrise (a late event) is also a likely synchronizer for risetimes; (3) the null hypothesis “risetimes distance to summer sunrise do not depend on *D*_*w*_” is rejected at this level: summer sunrise (an early event) is not a synchronizer for risetimes. Notice that second and third null hypotheses can also tested by contrasting {*y_i_ ∓βx_i_*} against {*x_i_*}. Probabilistic values and Pearson’s coefficients of these tests are also provided in Table IV. These results are visualized in Figure 2 by the one-hour width boxing technique.^1^

A final statistical test was performed: a set of *N* = 8 normally distributed values with standard deviation equal to sample standard deviation was tested against {*x_i_*}. This random set simulates noon synchronization plus normal noise. The process was repeated *M* = 10^5^ times and slope values *X* were collected. Only 4.2 % of the simulations hit *X* lower than observed *m*_*s*_ for risetimes. Therefore a random process synced to noon is relative unlikely to reproduce the observed distribution of risetimes.^2^

The analysis can be mimicked for bedtimes and wakeful noon times with similar results; the only remarkable difference being noon unlikely synchronizes by alone wakeful noon times: non-zero *m*_*s*_ is statistically significant. The regularity in *m*_*s*_ values for all three timemarks should be expected from the one-to-one correspondence shown in Figure 1 (left panel).

Including hunter/gatherer data helps increasing observed values to *N* = 11 and helps extending latitude equatorward. The range of the predictor increases to almost five hours which makes the range of winter sunrise time increase to two and a half hours. This helps deciding the competition: observed values for hunter/gatherer societies visually fit in the slanted box, well apart from the vertical box. Notice that this is not an extrapolation of NTUS regression analysis but a description of data placement which can easily be related to a univariate property: the variability of the risetimes. Table IV shows (first column) that NTUS risetime variability equals 2*s* = 46 min and increases by 250 % to 2*s* = 112 min if hunter/gatherer data are included (see second column). In this computation risetimes were cast as a distance to solar noon. If they are measured as a distance to winter sunrise, variability would remain unmodified: Table V lists 2*s* = 51 min for the eleven observations.

Multiple linear regression analysis gives us the quantitative answer for this observation. Second column in Table IV shows noon is rejected as a likely synchronizer for every of the timemarks: non-zero *m*_*s*_ is statistically significant. Observed slope values for risetimes and wakeful noon times hit *m_s_ ∼ -β* in accordance with the late event.

The preceding discussion does not discriminate which of the late events —winter sunrise or summer sunset— trigger human behavior. The phase or placement helps answering this question. Figure 2 shows people abhor rising at night, well before sunrise. Therefore the latest sunrise, winter sunrise, arises as a hallmark for human activity. Table V lists multiple linear regression data where the response is cast as either time distance to winter sunrise or winter solar elevation angle at risetimes. In average, risetimes occur at *z* = −8.7(35)°, within the winter nautical twilight or only −0.78(31) h before winter sunrise. Notice that winter solar elevation angle at risetimes versus *D*_*w*_ hits *p > α* and thus the null hypothesis sustains at the standard level. It is the only occurrence when *z* is the response.

Figure 2 shows people abhor anticipating activity to sunrise whereas they find accommodation in extending their activities after sunset, specially modern leisure activities. This asymmetry is nowadays related to the influence of artificial light in human behavior. However artificial light equally operates before sunrise when people make scarce use of it. Therefore physiology should be playing some role. Circadian regulation is sensitive to natural light and can set alertness when it is present. On the contrary it takes some time to trigger sleepiness after sunset, the process which may now be influenced by artificial light. A question to address is to what extend these processes were asymmetrical in ancient times.

A preliminary answer to this was noted by Yetish *et al.*[12]: pre-industrial societies “typically awaken before sunrise”, that is close to sunrise, some one hour of advance in winter. But they “go to bed several hours after sunset” (*δt_s_∼*3.5 h of delay in winter). Even in this natural case the asymmetry is perceptible.

We can figure out two processes for sleep onset. One is driven by a photoreceptive mechanism and could be expressed by the time distance from winter sunset to bedtime *δt_s_*. Assuming that the pre-industrial, Tropical value propagate across latitude then bedtimes at mid-latitudes would follow winter sunset trend (see light dash-dot-dot strip band in Figure 2, labeled “photoreceptive”). They would occur some three to four hours earlier than observed values. If that were the case in ancient time bedtimes would have followed the opposite trend to risetimes. This should have lead to bimodal or interrupted sleep[23] which should have been more frequent poleward. This sleep pattern would be a way of achieve synchronization overturning: the next risetime must have followed the opposite trend to that of preceding bedtime. Interestingly enough, present dinner time in Western Europe changes with latitude following winter sunset trend and mostly occur 3 h after winter sunset; that is immediately before the photoreceptive synchronizer shown in Figure 2. Synchronization overturning occur now in the evening when ambient light conditions change through seasons.[13]

The other process is driven by the homeostatic sleep pressure (Process S), the propensity to sleep onset related to previous awake. This can be described by *T* + *t*_1_ *-t*_3_. In this case risetime trend should propagate through the sleep-wake cycle as observed in Figure 2 where the homeostatic sleep onset differs 8.7 h from next risetime. As a result, bedtimes increasingly delay from sunset: up to 7 h at 50° latitude in winter, compared to *δt_s_* = 3.5 h. Wakeful noon times also delay with latitude and locate at 3pm at mid-latitudes, compared to 1pm in pre-industrial, equatorial societies.

As noted by Yetish *et al.*[12] the disappearance of Western European bimodal sleep in favor of modern unimodal sleep meant a return to the pattern of pre-industrial, equatorial societies but it should be stressed that the pattern is not synced to noon. Instead, the process was synced to the light/dark cycle by the winter sunrise: the ephemerides which seems able to trigger human activity in modern societies. That way, wakefulness after sunset and under artificial light increased as much as the sleep homeostatic pressure could provide; on the contrary, risetimes were not advanced; nor an equal combination of both processes was observed.

As a result sleep-wake timemarks are “delayed” with increasing latitude if they are measured relative to solar noon. This delay only finds relief in summer, when subsolar point travels back toward observer and turns sunrise/sunset trends upside-down. Figure 4 shows sleep/wake and light/dark cycles in summer. Light/dark seasonality is evident comparing Figure 4 to Figure 2, human activity requires further explanation.

**Figure 4.**
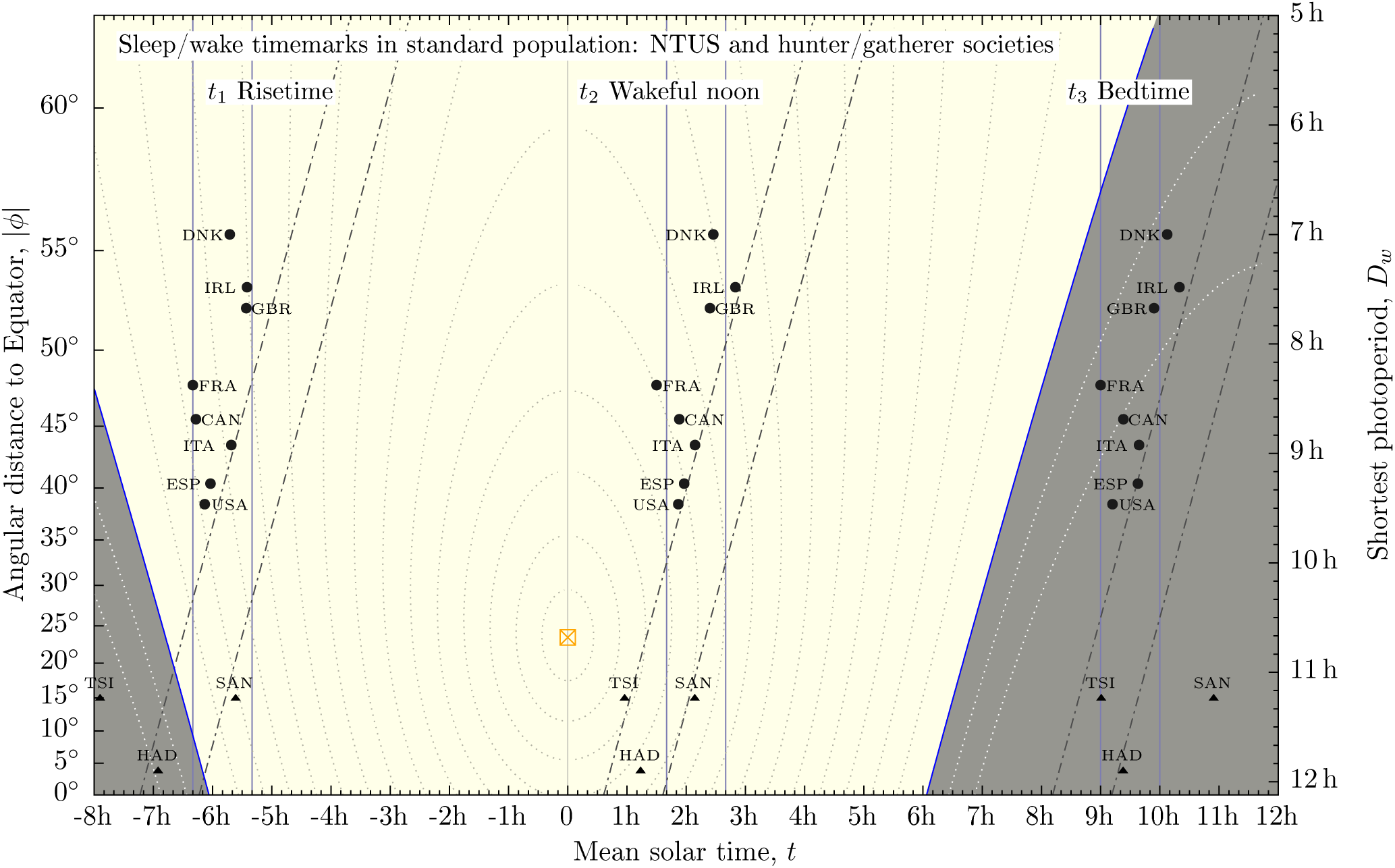
Timemarks extracted from standard population mean annual sleep-wake cycle in NTUS (circles) and hunter/gatherer pre-industrial societies from Table S2 in Yetish *et al.*[12] (triangles) against shortest photoperiod (right axis) or latitude (left axis). Blue solid lines show summer terminator (*z* = −0.83°). Dotted lines show solar elevation angle starting at *z* = −12°(outermost lines) in steps of 6°. Subsolar point (*z* = 90°) is shown by a crossed square. Lightest background shows summerphotoperiod; darkest background is summer night. Vertical solid lines box NTUS data in summer. Slanted dash-dotted lines are those of Figure 2. They visualize the effect of daylight saving time on NTUS data: sleep/wake cycle is advanced by one hour. Geographical data are listed in Table I; timemarks appear in Table II without DST for NTUS data.

Prior to 19th century it was seasonally adjusted, including labor starting times, which followed earlier sunrises. Hereafter, stable year round time schedules gained preference, which onset the winter sunrise time as a trigger year round. Since the first quarter of the 20th century daylight saving time (DST) remains as the main, if not unique, source of seasonality in human behavior of industrialized societies. It helps stabilizing social local time schedules, specially risetimes, by promoting one hour of advance in social solar time schedules. In figure 4 all NTUS values have been shifted one hour to the left following DST, while hunter/gatherer values are those of summer (see Table II). Slanted boxes keep the position shown in figure 2 and help visualizing data displacement. Summer daily rhythm in industrial societies in the mid-latitude range finds then more resemblance to hunter/gatherer, pre-industrial equatorial societies. It is perhaps for this reason that DST has been viewed as an equatorward travel[24] of human activity. However this is not anti-natural: either by seasonal clock adjustment (DST) or by seasonal time schedule adjustment human activity challenges the poleward travel of the subsolar point, which softens differences from Tropics to mid-latitudes. This can be noted in several ways when comparing Figure 2 and 4. First and most, subsolar point —the pole of the light hemisphere— can be observed in Figure 4 but not in Figure 2. It is relatively close to both Tropical regions and mid-latitudes, which makes life during the photoperiod more alike. For instance the number of solar elevation lines crossed from Hadza people’s to Danish people’s wakeful noon times dramatically changes: ten lines (Δ*z ∼* 60°) in winter; only three (Δ*z ∼* 18°) in summer.

Table VI lists multiple linear regression analysis for NTUS data with daylight saving times and hunter/gatherer pre-industrial societies in summer. For risetimes and wakeful noon times slope decreases to *m_s_ ∼ -β/*2 halfway from the late and noon events and confidence interval barely rejects events. Contrastingly bedtimes find *m_s_ ∼ -β/*10 and confidence interval sustains meridional synchronization in full accordance to what is observed in Figure 4. The difference in slope values is related to the fact that in summer *E*_hg_(*T* + *t*_1_ *- t*_3_) = 7.4 h (see Table II), one hour smaller than NTUS value.

### IV.2. Sleep and other personal care cycle: mid-latitudes to pole

HETUS data helps extending the range of the predictor in the opposite direction: poleward. However it is the “sleep and other personal care” that can now be analyzed. Figure 3 visualizes that societies above 54° advances their sleep/wake cycle relative to the winter sunrise. In other words, winter sunrise is so delayed that it no longer helps as a synchronizer. The corresponding advance of winter sunset and, in general, the large decrease of the photoperiod also helps breaking the trend of the winter sunrise.

As a consequence Table VII shows noon as the only event which sustains synchronization in the range 36° to 61°. Slope values lie in the range of *β/*10 in this case, smaller than results in Table IV.

Notwithstanding this “sleep and other personal care” data in the range of mid-latitudes (36° to 54°, see Table VIII) still reproduces results of the sleep-wake cycle (see Table IV) with increased number of observations. Noon and late events play similar roles and slope values in the range of *m_s_ ∼ -β/*2, not far from the value quoted in Table IV for NTUS data. It should be mentioned that the number of observations increases to *N* = 16 or *N* = 12 depending on whether duplicate values for Spain, Italy, France and Great Britain enter in the analysis or not.

Finally Table IX shows multiple linear regression results above 54°. Sample size (*N* = 7) is relatively small and above 54° the gradient *dD_w_/dφ* (see Equation (1)) is exceedingly large, while winter twilight lines open wide as their gradient decreases. All these issues jeopardize contrasting human activity to the light/dark cycle. This is noted by large confidence intervals in this analysis.

## V CONCLUSIONS

Below 54° latitude winter sunrise —the latest year round, the beginning to the region of permanent photoperiod— triggers human activity and is a relevant synchronizer for risetimes. Above that level risetimes must be advanced relative to winter sunrise because, which is exceedingly delayed and winter photoperiod is exceedingly short. In so doing, societies in this range match a lark behavior —risetime significantly advanced to sunrise— which is mostly abhorred elsewhere.

Also globally speaking sleep onset seems to be driven by the homeostatic process in a way that winter sunrise trend is also observed at bedtimes. Either people is tired or they are forecasting the following risetime.

Notices that this means bedtimes distance to noon increases as angular distance to Equator increases and they look delayed. Interestingly, this behavior increases wakefulness after sunset in winter, and promotes human activity under artificial light. That circumstance is not socially abhorred in the evening, whereas activity at dawn is mostly abhorred.

Seasonality —different behavior across seasons— is the only way this trend can be partially broken in summer so that mid-latitude societies return to the Equatorial pattern. In modern days this is mostly achieved by daylight saving time.

## ACKNOWLEDGMENTS

This work was not funded by any means. It was stoically endured by author’s wife, which codenamed it as “Canadian bidets”. No offense intended to the great Canadian people.

The author expresses his gratitude to the institutions which granted access to time use survey microdata.

## Appendix A: Geographical data and observed values

**Table I.** Overview of geographical data and ephemerides for participating countries. Population weighted median latitude is rounded to one tenth of degree. Photoperiod *D*_*w*_ displays the shortest photoperiod year round and it is a function of latitude and Earth’s obliquity (see Equation (1)). Winter sunrises and winter sunsets are given as a distance to solar noon in decimal hours. Time offset is the difference between solar noon and local time midday (rounded to one minute). Time offset and a factor *T/*2 = 12 h must be added to 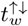 to get winter sunrise/sunset local times. Data from hunter/gatherer pre-industrial societies come from Yetish *et al.*[12]. No time-offset is needed in this case. For each subset sample average value, range and the variability expressed as twice sample standard deviation are listed. Photoperiod and latitude values appear in Figures 2, 3 and 4.

| Country/People | Label | Latitude<br>$\phi$ | Photoperiod<br>$D_w$ | Sunrise/set<br>$t_w^{\uparrow\downarrow} = \mp D_w/2$ | Time offset<sup>a</sup><br>$\delta$ |
| --- | --- | --- | --- | --- | --- |
| $\phi < 54^\circ$ | | | | | |
| United States | USA | 38.5° | 9.46 h | $\mp 4.73$ h | 8 min |
| Spain | ESP | 40.4° | 9.28 h | $\mp 4.64$ h | 72 min |
| Bulgaria | BGR | 42.7° | 9.02 h | $\mp 4.51$ h | 21 min |
| Italy | ITA | 43.6° | 8.92 h | $\mp 4.46$ h | 11 min |
| Canada | CAN | 45.5° | 8.69 h | $\mp 4.34$ h | 17 min |
| Slovenia | SVN | 46.1° | 8.62 h | $\mp 4.31$ h | 2 min |
| France | FRA | 47.8° | 8.38 h | $\mp 4.19$ h | 50 min |
| Belgium | BEL | 50.9° | 7.92 h | $\mp 3.96$ h | 42 min |
| Germany | DEU | 51.0° | 7.89 h | $\mp 3.94$ h | 23 min |
| Poland | POL | 51.8° | 7.77 h | $\mp 3.88$ h | −17 min |
| United Kingdom | GBR | 52.3° | 7.67 h | $\mp 3.84$ h | 6 min |
| Ireland | IRL | 53.3° | 7.48 h | $\mp 3.74$ h | 25 min |
| Average | $E(\{x_i\})$ | 47.0° | 8.43 h | $\mp 4.21$ h | 22 min |
| Range | $\Delta\{x_i\}_{\min}^{\max}$ | 14.8° | 119 min | 59 min | 89 min |
| Variability | $2s(\{x_i\})$ | 9.9° | 80 min | 40 min | 48 min |
| $\phi > 54^\circ$ | | | | | |
| Lithuania | LIT | 54.9° | 7.17 h | $\mp 3.58$ h | 24 min |
| Denmark | DNK | 55.7° | 7.00 h | $\mp 3.50$ h | 13 min |
| Latvia | LVA | 56.9° | 6.70 h | $\mp 3.35$ h | 24 min |
| Sweden | SWE | 59.2° | 6.08 h | $\mp 3.04$ h | −5 min |
| Estonia | EST | 59.4° | 6.03 h | $\mp 3.02$ h | 21 min |
| Norway | NOR | 59.9° | 5.87 h | $\mp 2.94$ h | 18 min |
| Finland | FIN | 61.1° | 5.45 h | $\mp 2.72$ h | 20 min |
| Average | $E(\{x_i\})$ | 58.2° | 6.33 h | $\mp 3.16$ h | 16 min |
| Range | $\Delta\{x_i\}_{\min}^{\max}$ | 6.2° | 103 min | 52 min | 29 min |
| Variability | $2s(\{x_i\})$ | 4.7° | 76 min | 38 min | 20 min |
| *Both combined* |  |  |  |  |  |
| Average | $E(\{x_i\})$ | 51.1° | 7.65 h | $\mp 3.83$ h | 20 min |
| Range | $\Delta\{x_i\}_{\min}^{\max}$ | 22.6° | 241 min | 121 min | 89 min |
| Variability | $2s(\{x_i\})$ | 13.8° | 146 min | 73 min | 39 min |
| *Hunter/gatherer* |  |  |  |  |  |
| Hadza | HAD | −3.7° | 11.90 h | $\mp 5.95$ h | |
| Tsimane | TSI | −14.9° | 11.24 h | $\mp 5.62$ h | |
| San | SAN | −14.9° | 11.24 h | $\mp 5.62$ h | |
| Average | $E(\{x_i\})$ | −11.2° | 11.46 h | $\mp 5.73$ h | |
| Range | $\Delta\{x_i\}_{\min}^{\max}$ | 11.2° | 40 min | 20 min | |
| Variability | $2s(\{x_i\})$ | 12.9° | 46 min | 23 min | |

<sup>a</sup> During daylight saving time $\delta$ must be increased in 60 min

**Table II.** Timemarks obtained from sleep/wake cycle in national time use surveys (NTUS) and in hunter/gatherer pre-industrial societies from Yetish *et al.*[12]. Timemarks are all expressed as a distance to solar noon in decimal hours. NTUS data are valid during winter time; one hour must be subtracted during summer time. Hadza people exhibit no seasonality since their distance to Equator is negligible small (see Table I). Winter solar elevation angle at risetime *z*_*w*_ and summer solar elevation angle at bedtimes *z*_*s*_ are also provided. They show the lowest and largest possible values year round, either case the worst case scenario for rising or sleep onset. Distance computes the difference from bedtime to next risetime. Values are displayed in Figure 1 (left), 2 and 4.

| Sleep timing in industrial and pre-industrial societies |  |  |  |  |  |  |  |
| --- | --- | --- | --- | --- | --- | --- | --- |
| Country/People | Label | Risetime |  | Wakeful noon | Bedtime |  | Bedtime to risetime |
| | | $t_1$ | $z_w$ | $t_2$ | $t_3$ | $z_s$ | $T + t_1 - t_3$ |
| Industrial/NTUS |  |  |  |  |  |  |  |
| United States | USA | -5.1 h | -5° | 2.9 h | 10.2 h | -17° | 8.7 h |
| Spain | ESP | -5.2 h | -7° | 3.0 h | 10.6 h | -18° | 8.2 h |
| Italy | ITA | -4.7 h | -3° | 3.1 h | 10.6 h | -16° | 8.7 h |
| Canada | CAN | -5.3 h | -9° | 2.9 h | 10.4 h | -12° | 8.3 h |
| France | FRA | -5.3 h | -11° | 2.7 h | 10.0 h | -8° | 8.7 h |
| United Kingdom | GBR | -4.4 h | -5° | 3.4 h | 10.9 h | -9° | 8.7 h |
| Ireland | IRL | -4.4 h | -6° | 3.8 h | 11.3 h | -10° | 8.2 h |
| Denmark | DNK | -4.7 h | -9° | 3.5 h | 11.0 h | -7° | 8.3 h |
| Average | $E(\{x_i\})$ | -4.9 h | -7° | 3.2 h | 10.6 h | -12° | 8.5 h |
| Range | $\Delta\{x_i\}_{\min}^{\max}$ | 55 min | 8° | 70 min | 80 min | 11° | 30 min |
| Variability | $2s(\{x_i\})$ | 46 min | 5° | 46 min | 52 min | 8° | 26 min |
| Pre-industrial/Hunter-gatherer (winter) |  |  |  |  |  |  |  |
| †Hadza | HAD | -6.9 h | -14° | 1.2 h | 9.4 h |  | 7.7 h |
| ‡Chico Mendes | CME | -6.8 h | -15° | 1.4 h | 9.5 h |  | 7.7 h |
| ‡Chico Mendes | CMA | -6.9 h | -17° | 1.0 h | 8.8 h |  | 8.2 h |
| †Tsimane | TSI | -7.2 h | -22° | 0.8 h | 8.7 h |  | 8.1 h |
| †San | SAN | -5.9 h | -5° | 1.9 h | 9.7 h |  | 8.4 h |
| *Toba/Qom | TQE | -5.2 h | 1° | 2.2 h | 9.6 h |  | 9.2 h |
| *Toba/Qom | TQN | -5.4 h | -1° | 1.6 h | 8.6 h |  | 10.0 h |
| Average | $E(\{x_i\})$ | -6.3 h | -11° | 1.4 h | 9.2 h | | 8.5 h |
| Range | $\Delta\{x_i\}_{\min}^{\max}$ | 120 min | 23° | 87 min | 63 min | | 138 min |
| Variability | $2s(\{x_i\})$ | 99 min | 17° | 62 min | 54 min | | 101 min |
| Pre-industrial/Hunter-gatherer (summer) |  |  |  |  |  |  |  |
| †Hadza | HAD | -6.9 h |  | 1.2 h | 9.4 h | -43° | 7.7 h |
| ‡Chico Mendes | CME | -6.8 h |  | 1.4 h | 9.5 h | -40° | 7.7 h |
| ‡Chico Mendes | CMA | -6.9 h |  | 1.0 h | 8.8 h | -33° | 8.2 h |
| †Tsimane | TSI | -7.9 h |  | 0.6 h | 9.0 h | -32° | 7.1 h |
| †San | SAN | -5.6 h |  | 2.7 h | 10.9 h | -48° | 7.5 h |
| *Toba/Qom | TQE | -6.2 h |  | 1.8 h | 9.9 h | -34° | 7.9 h |
| *Toba/Qom | TQN | -6.3 h |  | 1.2 h | 8.7 h | -22° | 9.0 h |
| Average | $E(\{x_i\})$ | -6.7 h | | 1.4 h | 9.5 h | -36° | 7.9 h |
| Range | $\Delta\{x_i\}_{\min}^{\max}$ | 137 min | | 126 min | 134 min | 26° | 115 min |
| Variability | $2s(\{x_i\})$ | 86 min | | 81 min | 91 min | 17° | 73 min |

**Table III.** Timemarks obtained from “sleep and other personal care” cycle in NTUS (starred) and HETUS (non-starred) subsets. Size refers to the number of respondents of each survey. Timemarks are all expressed as a distance to solar noon in decimal hours. They list winter time values; one hour must be subtracted during summer time. Winter solar elevation angle at risetime *z*_*w*_ and summer solar elevation angle at bedtimes *z*_*s*_ are also provided. They show the lowest and largest possible values year round, either case the worst case scenario for rising or sleep onset. Distance computes the difference from bedtimes to next risetime. Values are displayed in Figure 1 (right) and 3.

| “Sleep and other personal care” timemarks |  |  |  |  |  |  |  |  |
| --- | --- | --- | --- | --- | --- | --- | --- | --- |
| Country | Label | Respondents | Risetime |  | Wakeful noon | Bedtime |  | Distance |
| | | | $t_1$ | $z_w$ | $t_2$ | $t_3$ | $z_s$ | $T + t_1 - t_3$ |
| $\phi < 54^\circ$ | | | | | | | | |
| *United States | USA | 131 914 | −4.6 h | 0° | 2.9 h | 10.2 h | −17° | 9.2 h |
| Spain | ESP | 37 512 | −4.7 h | −1° | 3.0 h | 10.6 h | −18° | 8.7 h |
| *Spain | ESP | 16 571 | −4.7 h | −1° | 3.0 h | 10.5 h | −17° | 8.8 h |
| Bulgaria | BGR | 11 506 | −5.0 h | −6° | 2.7 h | 10.1 h | −13° | 8.8 h |
| *Italy | ITA | 32 068 | −4.4 h | 0° | 3.3 h | 10.6 h | −16° | 9.0 h |
| Italy | ITA | 38 149 | −4.5 h | −1° | 3.3 h | 10.8 h | −16° | 8.7 h |
| *Canada | CAN | 17 107 | −4.8 h | −5° | 3.1 h | 10.2 h | −11° | 9.0 h |
| Slovenia | SVN | 10 186 | −5.0 h | −7° | 2.6 h | 10.0 h | −9° | 9.0 h |
| *France | FRA | 25 617 | −4.8 h | −6° | 2.8 h | 10.0 h | −8° | 9.2 h |
| France | FRA | 12 913 | −5.0 h | −8° | 2.5 h | 10.0 h | −8° | 9.0 h |
| Belgium | BEL | 10 676 | −4.7 h | −7° | 2.8 h | 10.3 h | −7° | 9.0 h |
| Germany | DEU | 22 597 | −4.9 h | −8° | 2.8 h | 10.4 h | −8° | 8.7 h |
| Poland | POL | 35 212 | −4.4 h | −5° | 3.1 h | 10.4 h | −8° | 9.2 h |
| *United Kingdom | GBR | 15 409 | −4.1 h | −3° | 3.6 h | 10.7 h | −9° | 9.2 h |
| United Kingdom | GBR | 14 810 | −4.3 h | −4° | 3.4 h | 10.9 h | −9° | 8.8 h |
| *Ireland | IRL | 1834 | −3.9 h | −2° | 3.8 h | 11.1 h | −9° | 9.0 h |
| $\phi > 54^\circ$ | | | | | | | | |
| *Denmark | DNK | 5833 | −4.4 h | −7° | 3.5 h | 10.6 h | −5° | 9.0 h |
| Lithuania | LIT | 7418 | −5.2 h | −13° | 2.3 h | 9.6 h | 0° | 9.2 h |
| Latvia | LVA | 5786 | −5.1 h | −12° | 2.6 h | 10.1 h | −2° | 8.8 h |
| Sweden | SWE | 7612 | −4.4 h | −9° | 3.4 h | 11.1 h | −4° | 8.5 h |
| Estonia | EST | 8899 | −4.8 h | −12° | 2.8 h | 10.5 h | −1° | 8.7 h |
| Norway | NOR | 5458 | −4.5 h | −10° | 3.4 h | 11.0 h | −3° | 8.5 h |
| Finland | FIN | 8253 | −4.8 h | −13° | 2.8 h | 10.5 h | 0° | 8.7 h |

## Appendix B: Multiple linear regression analysis results

### 1. Sleep/wake cycle

**Table IV.** Multiple linear regression analysis for sleep-wake cycle timemarks (winter values for hunter/gather and winter time for NTUS). Each analysis takes the winter photoperiod as the predictor {*x_i_*} and the response {*y_i_*}is distance to noon. Probabilistic values and Pearson’s coefficient are also given for distance to late event and distance to early event. They are obtained by testing {*y_i_ ∓ βx_i_*} against {*x_i_*}. Predictor and responses are displayed in Figure 2.

| Regression analysis for sleep/wake cycle timemarks |  |  |  |
| --- | --- | --- | --- |
|  |  | NTUS | NTUS and preindustrial |
| **Predictor is winter photoperiod, $x = D_w$** |  |  |  |
| Observations | $N$ | 8 | 13 |
| Range | $\Delta\{x_i\}_{\min}^{\max}$ | 148 min | 294 min |
| Variability | $2s(\{x_i\})$ | 54 min | 96 min |
| Average | $E(\{x_i\})$ | 8.36 h | 9.43 h |
| **Response is risetimes distance to solar noon, $y = t_1$** |  |  |  |
| Range | $\Delta\{y_i\}_{\min}^{\max}$ | 55 min | 167 min |
| Variability | $2s(\{y_i\})$ | 46 min | 103 min |
| Average | $E(y_i)$ | -4.90(27) h | -5.37(31) h |
| Slope: value | $m_s$ | -17(19) min h<sup>-1</sup> | -27(12) min h<sup>-1</sup> |
| Slope: confidence interval | $CI$ | [-1.20, +0.09]$\beta$ | [-1.30, -0.48]$\beta$ |
| Cumulative probability for $m_s$ $P(X < m_s)$ | | 4.3 % | 0.2 % |
| The null hypothesis sustains for event |  | Noon/Late | Late |
| $p$-value | $p$ | 0.08/0.14 | 0.56 |
| Pearson's coefficient | $R^2$ | 0.42/0.32 | 0.032 |
| The null hypothesis is overruled for event |  | Early | Noon/Early |
| $p$-value | $p$ | $< 10^{-3}$ | $< 10^{-3}/< 10^{-7}$ |
| Pearson's coefficient | $R^2$ | 0.87 | 0.68/0.94 |
| **Response is wakeful noon times distance to solar noon, $y = t_2$** |  |  |  |
| Range | $\Delta\{y_i\}_{\min}^{\max}$ | 70 min | 185 min |
| Variability | $2s(\{y_i\})$ | 46 min | 111 min |
| Average | $E(y_i)$ | 3.15(25) h | 2.53(24) h |
| Slope: value | $m_s$ | -19(18) min h<sup>-1</sup> | -31.9(94) min h<sup>-1</sup> |
| Slope: confidence interval | $CI$ | [-1.22, -0.05]$\beta$ | [-1.38, -0.75]$\beta$ |
| Cumulative probability for $m_s$ $P(X < m_s)$ | | 2.7 % | 0.1 % |
| The null hypothesis sustains for event |  | Late | Late |
| $p$-value | $p$ | 0.18 | 0.66 |
| Pearson's coefficient | $R^2$ | 0.28 | 0.018 |
| The null hypothesis is overruled for event |  | Noon/Early | Noon/Early |
| $p$-value | $p$ | 0.038/$< 10^{-3}$ | $< 10^{-4}/< 10^{-7}$ |
| Pearson's coefficient | $R^2$ | 0.54/0.89 | 0.84/0.95 |
| **Response is bedtimes distance to solar noon, $y = t_3$** |  |  |  |
| Range | $\Delta\{y_i\}_{\min}^{\max}$ | 80 min | 162 min |
| Variability | $2s(\{y_i\})$ | 52 min | 101 min |
| Average | $E(y_i)$ | 10.63(30) h | 10.08(28) h |
| Slope: value | $m_s$ | -19(22) min h<sup>-1</sup> | -27(11) min h<sup>-1</sup> |
| Slope: confidence interval | $CI$ | [-1.37, +0.08]$\beta$ | [-1.27, -0.54]$\beta$ |
| Cumulative probability for $m_s$ $P(X < m_s)$ | | 3.9 % | 0.2 % |
| The null hypothesis sustains for event |  | Noon/Late | Late |
| $p$-value | $p$ | 0.072/0.28 | 0.58 |
| Pearson's coefficient | $R^2$ | 0.44/0.19 | 0.028 |
| The null hypothesis is overruled for event |  | Early | Noon/Early |
| $p$-value | $p$ | $< 10^{-2}$ | $< 10^{-3}/< 10^{-6}$ |
| Pearson's coefficient | $R^2$ | 0.84 | 0.73/0.92 |

**Table V.**
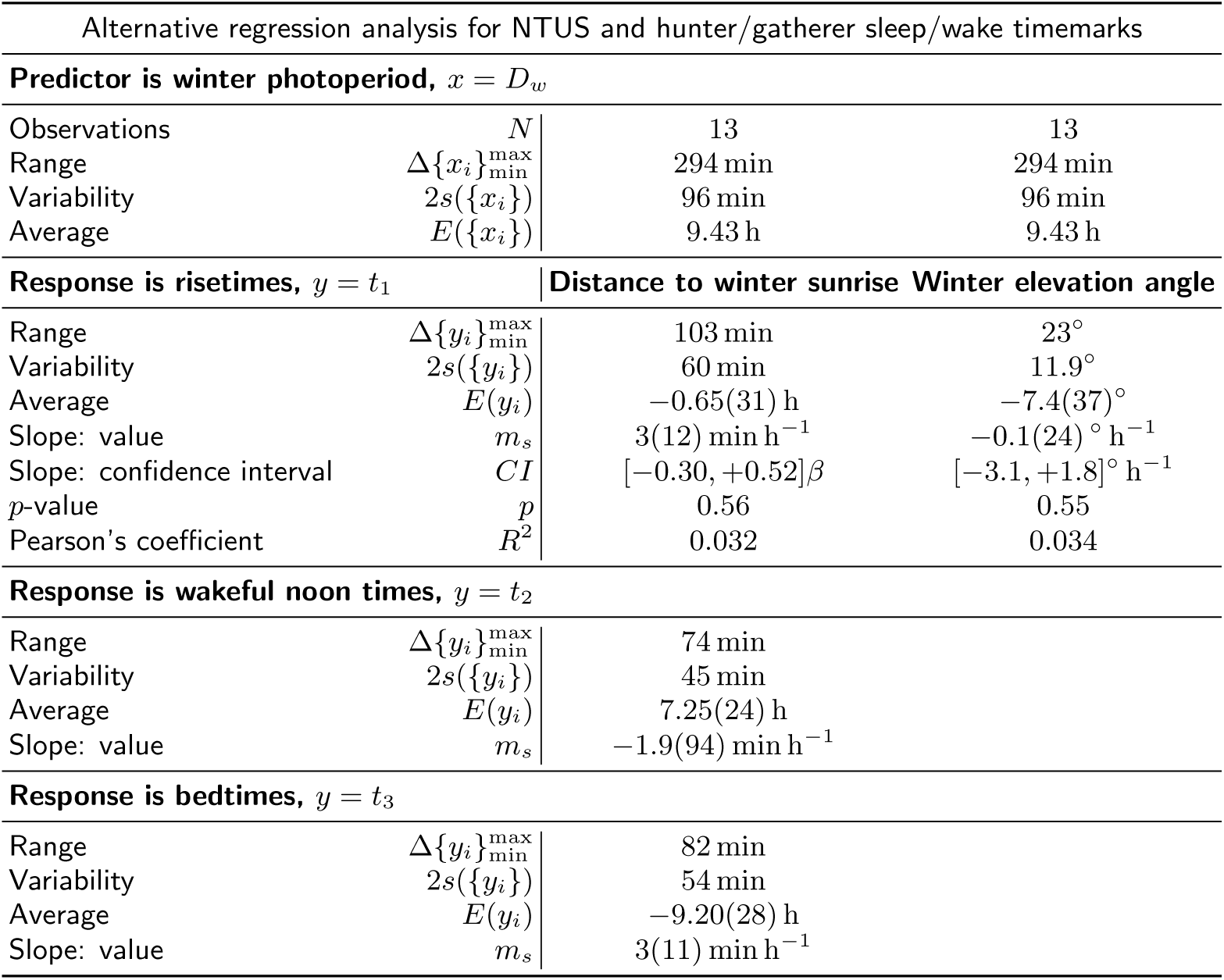
Alternative multiple linear regression analysis for the sleep/wake timemarks in NTUS and hunter/gatherer pre-industrial societies (winter values for hunter/gather and winter time for NTUS). Each analysis takes the winter photoperiod as the predictor {*x_i_*} and the response {*y_i_*}is distance to winter sunrise (first column) and winter solar elevation angle (second column). Notice the decrease in the variability of distance to winter sunrise compared to distance to noon (see Table IV). Also notice average response is close to winter terminator: three quarters of an hour in advance and within the nautical twilight. Average value for bedtimes represent distance to the following winter sunrise time. Predictor and responses are displayed in Figure 2.

**Table VI.** Multiple linear regression analysis for the sleep/wake timemarks in NTUS and hunter/gatherer pre-industrial societies in summer. NTUS data (see Table IV) are corrected by DST and advanced by one whole hour. Each analysis takes the winter photoperiod as the predictor {*x_i_*}and the response {*y_i_*}is is distance to noon. Predictor and responses are displayed in Figure 4.

| Regression analysis for NTUS data with DST and hunter-gatherer summer sleep/wake timemarks |  |  |
| --- | --- | --- |
|  |  | DST |
| **Predictor is winter photoperiod, $x = D_w$** |  |  |
| Observations | $N$ | 13 |
| Range | $\Delta\{x_i\}_{\min}^{\max}$ | 294 min |
| Variability | $2s(\{x_i\})$ | 96 min |
| Average | $E(\{x_i\})$ | 9.43 h |
| **Response is risetimes, $y = t_1$** |  |  |
| Range | $\Delta\{y_i\}_{\min}^{\max}$ | 107 min |
| Variability | $2s(\{y_i\})$ | 63 min |
| Average | $E(y_i)$ | -6.14(23) h |
| Slope: value | $m_s$ | -14.6(89) min h<sup>-1</sup> |
| Slope: confidence interval | $CI$ | $[-0.78, -0.19]\beta$ |
| Cumulative probability for $m_s$ $P(X < m_s)$ | | 0.5 % |
| $p$-value | $p$ | $< 10^{-2}$ |
| Pearson's coefficient | $R^2$ | 0.54 |
| **Response is wakeful noon times, $y = t_2$** |  |  |
| Range | $\Delta\{y_i\}_{\min}^{\max}$ | 132 min |
| Variability | $2s(\{y_i\})$ | 79 min |
| Average | $E(y_i)$ | 1.76(25) h |
| Slope: value | $m_s$ | -19.9(97) min h<sup>-1</sup> |
| Slope: confidence interval | $CI$ | $[-0.99, -0.34]\beta$ |
| Cumulative probability for $m_s$ $P(X < m_s)$ | | 0.3 % |
| $p$-value | $p$ | $< 10^{-3}$ |
| Pearson's coefficient | $R^2$ | 0.65 |
| **Response is bedtimes, $y = t_3$** |  |  |
| Range | $\Delta\{y_i\}_{\min}^{\max}$ | 162 min |
| Variability | $2s(\{y_i\})$ | 84 min |
| Average | $E(y_i)$ | 9.31(37) h |
| Slope: value | $m_s$ | -15(14) min h<sup>-1</sup> |
| Slope: confidence interval | $CI$ | $[-0.99, -0.03]\beta$ |
| Cumulative probability for $m_s$ $P(X < m_s)$ | | 2.3 % |
| $p$-value | $p$ | 0.041 |
| Pearson's coefficient | $R^2$ | 0.33 |

## 2. Sleep and other personal care cycle

**Table VII.** Multiple linear regression analysis for “sleep and other personal care” timemarks. Each analysis takes the winter photoperiod as the predictor {*x_i_*}and the response {*y_i_*} is distance to noon. Probabilistic values and Pearson’s coefficient are also given for distance to late event and distance to early event. They are obtained by testing {*y_i_ ∓βx_i_*} against {*x_i_*}. Left column display analysis for all available data. In the right column HETUS duplicates were removed. Predictor and responses are displayed in Figure 3.

| Regression analysis for “Sleep and other personal care” cycle timemarks. Range: $\vert \phi\vert > 36^\circ$ | | | |
| --- | --- | --- | --- |
|  |  | All data | No duplicates |
| **Predictor is winter photoperiod, $x = D_w$** |  |  |  |
| Observations | $N$ | 23 | 19 |
| Range | $\Delta\{x_i\}_{\min}^{\max}$ | 241 min | 241 min |
| Variability | $2s(\{x_i\})$ | 71 min | 73 min |
| Average | $E(\{x_i\})$ | 7.81 h | 7.65 h |
| **Response is risetimes distance to solar noon, $y = t_1$** |  |  |  |
| Range | $\Delta\{y_i\}_{\min}^{\max}$ | 79 min | 79 min |
| Variability | $2s(\{y_i\})$ | 40 min | 41 min |
| Average | $E(y_i)$ | -4.66(15) h | -4.66(17) h |
| Slope: value | $m_s$ | -1.1(76) min h<sup>-1</sup> | -0.1(87) min h<sup>-1</sup> |
| Slope: confidence interval | $CI$ | $[-0.29, +0.22]\beta$ | $[-0.32, +0.26]\beta$ |
| Cumulative probability for $m_s$ $P(X < m_s)$ | | 38.2 % | 41.0 % |
| The null hypothesis sustains for event |  | Noon | Noon |
| $p$-value | $p$ | 0.77 | 0.83 |
| Pearson's coefficient | $R^2$ | $< 10^{-2}$ | $< 10^{-2}$ |
| The null hypothesis is overruled for event |  | Late/Early | Late/Early |
| $p$-value | $p$ | $< 10^{-7}/< 10^{-8}$ | $< 10^{-5}/< 10^{-6}$ |
| Pearson's coefficient | $R^2$ | 0.75/0.81 | 0.75/0.81 |
| **Response is wakeful noon times distance to solar noon, $y = t_2$** |  |  |  |
| Range | $\Delta\{y_i\}_{\min}^{\max}$ | 94 min | 94 min |
| Variability | $2s(\{y_i\})$ | 47 min | 47 min |
| Average | $E(y_i)$ | 3.01(17) h | 3.01(19) h |
| Slope: value | $m_s$ | -2.5(88) min h<sup>-1</sup> | -2.7(98) min h<sup>-1</sup> |
| Slope: confidence interval | $CI$ | $[-0.38, +0.21]\beta$ | $[-0.42, +0.24]\beta$ |
| Cumulative probability for $m_s$ $P(X < m_s)$ | | 27.9 % | 27.6 % |
| The null hypothesis sustains for event |  | Noon | Noon |
| $p$-value | $p$ | 0.57 | 0.57 |
| Pearson's coefficient | $R^2$ | 0.016 | 0.019 |
| The null hypothesis is overruled for event |  | Late/Early | Late/Early |
| $p$-value | $p$ | $< 10^{-5}/< 10^{-6}$ | $< 10^{-4}/< 10^{-5}$ |
| Pearson's coefficient | $R^2$ | 0.67/0.74 | 0.67/0.74 |
| **Response is bedtimes distance to solar noon, $y = t_3$** |  |  |  |
| Range | $\Delta\{y_i\}_{\min}^{\max}$ | 89 min | 89 min |
| Variability | $2s(\{y_i\})$ | 47 min | 47 min |
| Average | $E(y_i)$ | 10.45(17) h | 10.42(18) h |
| Slope: value | $m_s$ | -5.6(86) min h<sup>-1</sup> | -7.4(91) min h<sup>-1</sup> |
| Slope: confidence interval | $CI$ | $[-0.47, +0.10]\beta$ | $[-0.55, +0.06]\beta$ |
| Cumulative probability for $m_s$ $P(X < m_s)$ | | 9.4 % | 5.2 % |
| The null hypothesis sustains for event |  | Noon | Noon |
| $p$-value | $p$ | 0.19 | 0.11 |
| Pearson's coefficient | $R^2$ | 0.079 | 0.15 |
| The null hypothesis is overruled for event |  | Late/Early | Late/Early |
| $p$-value | $p$ | $< 10^{-5}/< 10^{-7}$ | $< 10^{-4}/< 10^{-6}$ |
| Pearson's coefficient | $R^2$ | 0.62/0.78 | 0.62/0.81 |

**Table VIII.** Multiple linear regression analysis for “sleep and other personal care” timemarks below 54°. Each analysis takes the winter photoperiod as the predictor {*x_i_* and the response {*y_i_*}is distance to noon. Probabilistic values and Pearson’s coefficient are also given for distance to late event and distance to early event. They are obtained by testing {*y_i_ ∓βx_i_*}against {*x_i_*}. Left column display analysis for all available data. In the right column HETUS duplicates were removed. Predictor and responses are displayed in Figure 3

| Regression analysis for “Sleep and other personal care” cycle timemarks. Range: $36^\circ < \vert \phi\vert < 54^\circ$ | | | |
| --- | --- | --- | --- |
|  |  | All data | No duplicates |
| **Predictor is winter photoperiod, $x = D_w$** |  |  |  |
| Observations | $N$ | 16 | 12 |
| Range | $\Delta\{x_i\}_{\min}^{\max}$ | 119 min | 119 min |
| Variability | $2s(\{x_i\})$ | 39 min | 40 min |
| Average | $E(\{x_i\})$ | 8.46 h | 8.43 h |
| **Response is risetimes distance to solar noon, $y = t_1$** |  |  |  |
| Range | $\Delta\{y_i\}_{\min}^{\max}$ | 67 min | 67 min |
| Variability | $2s(\{y_i\})$ | 40 min | 42 min |
| Average | $E(y_i)$ | −4.61(16) h | −4.61(21) h |
| Slope: value | $m_s$ | −15(15) min h<sup>−1</sup> | −16(19) min h<sup>−1</sup> |
| Slope: confidence interval | $CI$ | [−1.00, +0.02]$\beta$ | [−1.17, +0.11]$\beta$ |
| Cumulative probability for $m_s$ $P(X < m_s)$ | | 3.0 % | 4.6 % |
| The null hypothesis sustains for event |  | Noon/Late | Noon/Late |
| $p$-value | $p$ | 0.057/0.05 | 0.093/0.14 |
| Pearson's coefficient | $R^2$ | 0.24/0.25 | 0.26/0.21 |
| The null hypothesis is overruled for event |  | Early | Early |
| $p$-value | $p$ | $< 10^{-4}$ | $< 10^{-3}$ |
| Pearson's coefficient | $R^2$ | 0.76 | 0.76 |
| **Response is wakeful noon times distance to solar noon, $y = t_2$** |  |  |  |
| Range | $\Delta\{y_i\}_{\min}^{\max}$ | 80 min | 72 min |
| Variability | $2s(\{y_i\})$ | 44 min | 44 min |
| Average | $E(y_i)$ | 3.04(18) h | 3.03(22) h |
| Slope: value | $m_s$ | −14(17) min h<sup>−1</sup> | −16(20) min h<sup>−1</sup> |
| Slope: confidence interval | $CI$ | [−1.05, +0.12]$\beta$ | [−1.22, +0.14]$\beta$ |
| Cumulative probability for $m_s$ $P(X < m_s)$ | | 5.4 % | 5.2 % |
| The null hypothesis sustains for event |  | Noon/Late | Noon/Late |
| $p$-value | $p$ | 0.11/0.069 | 0.11/0.16 |
| Pearson's coefficient | $R^2$ | 0.17/0.22 | 0.24/0.19 |
| The null hypothesis is overruled for event |  | Early | Early |
| $p$-value | $p$ | $< 10^{-4}$ | $< 10^{-3}$ |
| Pearson's coefficient | $R^2$ | 0.68 | 0.72 |
| **Response is bedtimes distance to solar noon, $y = t_3$** |  |  |  |
| Range | $\Delta\{y_i\}_{\min}^{\max}$ | 67 min | 67 min |
| Variability | $2s(\{y_i\})$ | 41 min | 39 min |
| Average | $E(y_i)$ | 10.44(18) h | 10.39(19) h |
| Slope: value | $m_s$ | −11(17) min h<sup>−1</sup> | −14(18) min h<sup>−1</sup> |
| Slope: confidence interval | $CI$ | [−0.92, +0.20]$\beta$ | [−1.07, +0.11]$\beta$ |
| Cumulative probability for $m_s$ $P(X < m_s)$ | | 9.1 % | 4.8 % |
| The null hypothesis sustains for event |  | Noon | Noon/Late |
| $p$-value | $p$ | 0.19 | 0.098/0.076 |
| Pearson's coefficient | $R^2$ | 0.12 | 0.25/0.28 |
| The null hypothesis is overruled for event |  | Late/Early | Early |
| $p$-value | $p$ | 0.028/$< 10^{-3}$ | $< 10^{-3}$ |
| Pearson's coefficient | $R^2$ | 0.3/0.66 | 0.76 |

**Table IX.** Multiple linear regression analysis for “sleep and other personal care” timemarks above 54°. Each analysis takes the winter photoperiod as the predictor {*x_i_*} and the response {*y_i_*} is distance to noon. Probabilistic values and Pearson’s coefficient are also given for distance to late event and distance to early event. They are obtained by testing {*y_i_ ∓ βx_i_*} against {*x_i_*}. Predictor and responses are displayed in Figure 3

|  |  | All data |
| --- | --- | --- |
| **Predictor is winter photoperiod, $x = D_w$** |  |  |
| Observations | $N$ | 7 |
| Range | $\Delta\{x_i\}_{\min}^{\max}$ | 103 min |
| Variability | $2s(\{x_i\})$ | 38 min |
| Average | $E(\{x_i\})$ | 6.33 h |
| **Response is risetimes distance to solar noon, $y = t_1$** |  |  |
| Range | $\Delta\{y_i\}_{\min}^{\max}$ | 52 min |
| Variability | $2s(\{y_i\})$ | 40 min |
| Average | $E(y_i)$ | -4.75(34) h |
| Slope: value | $m_s$ | -9(35) min h<sup>-1</sup> |
| Slope: confidence interval | $CI$ | $[-1.47, +0.85]\beta$ |
| Cumulative probability for $m_s$ | $P(X < m_s)$ | 23.6 % |
| The null hypothesis sustains for event |  | Noon/Late |
| $p$-value | $p$ | 0.52/0.19 |
| Pearson's coefficient | $R^2$ | 0.086/0.32 |
| The null hypothesis is overruled for event |  | Early |
| $p$-value | $p$ | 0.022 |
| Pearson's coefficient | $R^2$ | 0.68 |
| **Response is wakeful noon times distance to solar noon, $y = t_2$** |  |  |
| Range | $\Delta\{y_i\}_{\min}^{\max}$ | 72 min |
| Variability | $2s(\{y_i\})$ | 55 min |
| Average | $E(y_i)$ | 2.96(47) h |
| Slope: value | $m_s$ | -13(47) min h<sup>-1</sup> |
| Slope: confidence interval | $CI$ | $[-2.01, +1.15]\beta$ |
| Cumulative probability for $m_s$ | $P(X < m_s)$ | 23.3 % |
| The null hypothesis sustains for event |  | Noon/Late/Early |
| $p$-value | $p$ | 0.52/0.4/0.068 |
| Pearson's coefficient | $R^2$ | 0.089/0.15/0.52 |
| The null hypothesis is overruled for event |  |  |
| $p$-value | $p$ | |
| Pearson's coefficient | $R^2$ | |
| **Response is bedtimes distance to solar noon, $y = t_3$** |  |  |
| Range | $\Delta\{y_i\}_{\min}^{\max}$ | 89 min |
| Variability | $2s(\{y_i\})$ | 62 min |
| Average | $E(y_i)$ | 10.49(44) h |
| Slope: value | $m_s$ | -30(44) min h<sup>-1</sup> |
| Slope: confidence interval | $CI$ | $[-2.47, +0.50]\beta$ |
| Cumulative probability for $m_s$ | $P(X < m_s)$ | 6.9 % |
| The null hypothesis sustains for event |  | Noon/Late |
| $p$-value | $p$ | 0.15/0.98 |
| Pearson's coefficient | $R^2$ | 0.37/< 10<sup>-3</sup> |
| The null hypothesis is overruled for event |  | Early |
| $p$-value | $p$ | 0.018 |
| Pearson's coefficient | $R^2$ | 0.7 |

## Footnotes

1 Notice that risetimes can also be placed in a slanted box with slope *m* = +*β* but box width must be much larger than one hour.

2 For instance testing the decimal fraction of the time offset *δ* against *D*_*w*_ results in a slope −2(16) min h^-1^, *p* = 0.78.

## REFERENCES

[1] A. A. Borbély, Human Neurobiology 1, 195 (1982).

[2] S. Daan, D. G. Beersma, and A. A. Borbély, The American journal of physiology 246, R161 (1984).

[3] A. C. Skeldon, A. J. K. Phillips, and D.-J. Dijk, Scientific Reports 7, 45158 (2017).

[4] A. A. Borbély, S. Daan, A. Wirz-Justice, and T. Deboer, Journal of Sleep Research 25, 131 (2016).

[5] T. Roenneberg, T. Kantermann, M. Juda, C. Vetter, and K. V. Allebrandt, Handbook of Experimental Pharmacology 217, 311 (2013).

[6] T. M. White and M. Terman, Psychiatry: Interpersonal and Biological Processes 742, 1193 (2003).

[7] T. Roenneberg, C. J. Kumar, and M. Merrow, Current Biology 17, R44 (2007).

[8] C. Randler, Chronobiology International 25, 1017 (2008).

[9] M. Miguel, V. C. de Oliveira, D. Pereira, and M. Pedrazzoli, Annals of Human Biology 41, 107 (2014).

[10] M. A. Leocadio-Miguel, F. M. Louzada, L. L. Duarte, R. P. Areas, M. Alam, M. V. Freire, J. Fontenele-Araujo, L. Menna-Barreto, and M. Pedrazzoli, Scientific Reports 7, 5437 (2017).

[11] C. Randler and A. Rahafar, Scientific Reports 7, 39976 (2017).

[12] G. Yetish, H. Kaplan, M. Gurven, B. Wood, H. Pontzer, P. R. Manger, C. Wilson, R. McGregor, and J. M. Siegel, Current biology: CB 25, 2862 (2015).

[13] J. M. Martín-Olalla, Scientific Reports 2018 8:1 8, 5350 (2018).

[14] Det Nationale Forkskningscenter for Velfærd, “Danish Time Use Survey: Danske Tidsanvendelseundersøgelsen,” Center for Survey and Survey/Register Data (distribuitor) (2001).

[15] Instituto Nacional de Estadística, “Spanish Time Use Survey: Encuesta de Empleo del Tiempo,” (2010).

[16] Bureau of Labor Statistics, “American Time Use Survey,” computer file (multi year data) (2012).

[17] L’Institut National de la Statisque et des Études Économiques, “French Time Use Survey. Enquête emploi du Temps et Décisions dans les couples,” (2010).

[18] Economic and Social Research Institute, “The Irish National Time-Use Survey,” Irish Social Sience Data Archive (distribuitor) (2005).

[19] L’Istituto nazionale di statistica (Istat), “Italian Time Use Survey: Uso del tempo,” (2009).

[20] Ipsos-RSL and Office of National Statistics, “United Kingdom Time Use Survey 2000 (computer file),” 3rd ed, Colchester, Essex: UK Data archive (distribuitor) (2003).

[21] Statistics Canada/Statisque Canada, “General Social Survey, Time Use. cycle 19,” computer file (2005).

[22] Statistics Finland and Statistics Sweden, “Harmonised European Time Use Survey [online database version 2.0],” (2005-2007).

[23] A. R. Ekirch, At day’s close: night in times past (Norton, 2005) p. 447.

[24] T. Kantermann, M. Juda, M. Merrow, and T. Roenneberg, Current Biology 17, 1996 (2007).

